# SHERLOC: An interpretable deep learning model for longitudinal circulating tumor DNA data in survival analysis

**DOI:** 10.64898/2026.06.04.730097

**Authors:** A. Mamann, J. Das, H. Benkirane, F. Bugiotti, E. Bernard, B. Besse, S. Michiels, P. H. Cournède

**Affiliations:** Université Paris-Saclay, Gustave Roussy, CentraleSupélec, INSERM, Cancer Data Science Unit, Villejuif, France; IHU PRISM National Precision Medicine Center in Oncology, Gustave Roussy, Villejuif, France; Université Paris-Saclay, CentraleSupélec, Lab of Mathematics and Computer Science (MICS), Gif-sur-Yvette, 91190, France; Université Paris-Saclay, CESP, INSERM U1018, labeled Ligue Contre le Cancer, Oncostat Team, Villejuif, France; Université Paris-Saclay, CNRS, CentraleSupélec, LISN, Gif-sur-Yvette, France; Gustave Roussy, Université Paris-Saclay, INSERM U981, Villejuif, France; Gustave Roussy, Université Paris-Saclay, Thoracic Group and International Center for Thoracic Cancers, Department of Cancer Medicine, Villejuif, France; Université Paris-Saclay, Faculty of Medicine, Paris, France; Gustave Roussy, Université Paris-Saclay, Office of Biostatistics and Epidemiology, Villejuif, France

## Abstract

Longitudinal circulating tumor DNA (ctDNA) measurements offer a noninvasive means to monitor treatment response, but clinical trial data present substantial methodological challenges due to high-dimensional short longitudinal ctDNA sequences and limited sample sizes. We introduce SHERLOC, a deep learning framework specifically designed for survival analysis using longitudinal on-treatment ctDNA data, which integrates shared temporal representations of gene-level variant allele frequencies, feature-specific temporal trajectories of panel-level ctDNA biomarkers, and survival-aware genomic representations pre-trained on a large pan-cancer tissue-biopsy dataset (MSK-CHORD), within an interpretable Cox proportional hazards framework. Benchmarked against diverse statistical, ensemble, and deep learning approaches in a non–small-cell lung cancer cohort from the phase III IMpower150 trial, SHERLOC consistently achieved superior survival discrimination and calibration, while remaining interpretable and robust to reductions in the number of available longitudinal liquid biopsy time points per patient. The resulting ctDNA-based risk score provided prognostic information both independent of and complementary to standard radiographic response assessments, and enabled patient stratification within homogeneous RECIST response groups—highlighting its potential as an early, non-invasive decision-support tool to guide treatment adaptation and patient management.

## Introduction

Imaging-based RECIST criteria have long been the standard for assessing treatment response and guiding clinical decision-making [1–3]; however, they remain imperfect surrogates of therapeutic efficacy [4, 5], as they provide an incomplete characterization of tumor biology, failing to capture critical molecular features such as genomic alterations and mutational burden. Longitudinal monitoring of circulating tumor DNA (ctDNA) has emerged as a minimally invasive approach to address these limitations through serial blood sampling [6, 7]. As the tumor-derived portion of cell-free DNA, ctDNA reflects the evolving molecular landscape of cancer. It has been shown to carry both prognostic and predictive information across multiple tumor types and treatment settings [8, 9], including insights not captured by conventional imaging assessments like RECIST [10, 11]. By enabling real-time monitoring of molecular responses, longitudinal ctDNA measurements offer a sensitive framework for early detection of therapeutic benefit or resistance. This has motivated efforts to standardize ctDNA-based response evaluation and to develop robust statistical and computational models for predicting patient outcomes [10–12]. Specifically, deep learning approaches are proving essential for integrating high-dimensional molecular features to advance precision medicine [13, 14].

Cell-free DNA (cfDNA) samples were collected at four predefined time points from 466 patients with non-small-cell lung cancer (NSCLC) enrolled in the randomized phase III IMpower150 clinical trial (NCT02366143) [10, 15]. IMpower150 evaluated the safety and efficacy of the anti–PD-L1 antibody atezolizumab in combina-tion with carboplatin and paclitaxel, with or without bevacizumab, compared with carboplatin, paclitaxel, and bevacizumab alone in chemotherapy-naïve patients with metastatic nonsquamous NSCLC. Baseline plasma was profiled using a 394-gene prototype of the FoundationOne® Liquid CDx assay (Foundation Medicine Inc.). Subsequently, on-treatment samples were evaluated via a custom fixed-panel assay specifically designed to monitor treatment-induced ctDNA dynamics [10].

Survival analyses were performed using a landmark approach, with overall survival (OS) defined from the time of the final on-treatment ctDNA collection. Patients were considered evaluable if plasma samples were available at baseline and at all subsequent on-treatment time points (Cycle 2 Day 1 [C2D1], C3D1, and C4D1), together with matched peripheral blood mononuclear cells (PBMCs). Patients who experienced a clinical event prior to the designated landmark were excluded from the analysis, resulting in landmark cohort of 386 patients at the C4D1 time point.

The data of interest consist of longitudinal variant allele frequencies (VAFs) measured in ctDNA for 109 genes—corresponding to genes mutated in at least five patients in the cohort—across the first four treatment cycles (BL, C2D1, C3D1, and C4D1). In addition, longitudinal panel-level cfDNA/ctDNA features (cfDNA concentration, number of driver mutations, mean VAF, and maximum VAF) were collected at the same time points, together with relevant clinical covariates including PD-L1 status, number of metastases, treatment arm, baseline tumor size, age, sex, and ECOG performance status.

This results in a high-dimensional longitudinal dataset characterized by more than 100 molecular dimensions but only a very small number of time points (4 timesteps). Such a data structure poses significant methodological challenges. Traditional approaches for longitudinal analysis, including classical statistical methods such as joint models, were not designed for this specific setting [16, 17]. Similarly, standard deep learning architectures for sequential data, such as recurrent neural networks (RNNs) or gated recurrent units (GRUs), typically require longer sequences to learn meaningful temporal dependencies and are therefore ill-adapted to short longitudinal trajectories [18, 19].

Importantly, increasing the number of time points through additional liquid biopsies is often not feasible in practice, as next-generation sequencing (NGS)-based profiling of cfDNA remains costly, especially when high sensitivity is needed for the detection of low ctDNA VAFs [20, 21]. Furthermore, the application of machine learning methods is constrained by the limited sample size inherent to clinical trials (e.g., 386 patients in our C4D1 landmark analysis), which restricts the complexity and number of parameters of models that can be reliably trained without overfitting [22, 23].

Finally, model interpretability represents a critical requirement in this clinical context. Beyond predictive performance, it is essential to quantify the contribution of individual molecular and clinical biomarkers to the final prediction, in order to support clinical decision-making and enable oncologists to interpret and trust model recommendations. These combined challenges motivate the development of novel methodological approaches tailored to high-dimensional, four-point longitudinal cfDNA profiles with limited sample sizes and strong interpretability constraints.

In this study, we introduce SHERLOC (Shared and Exclusive Representations of Longitudinal ctDNA data), a deep learning model for survival analysis based on longitudinal ctDNA measurements. SHERLOC leverages a hybrid survival modeling strategy tailored to short ctDNA trajectories. The model jointly captures (i) shared temporal patterns across gene-level variant allele frequencies (VAFs), (ii) feature-specific longitudinal dynamics of panel-level cfDNA biomarkers, and (iii) survival-informed genomic embeddings pre-trained on large external cohort. These components are integrated within an interpretable Cox proportional hazards layer, enabling both predictive performance and biological insight. We begin by detailing SHERLOC’s architecture and training protocol, followed by a comparative analysis against classical statistical methods, machine learning models, and alternative deep learning techniques. Performance is evaluated based on survival discrimination and calibration. We then demonstrate SHERLOC’s clinical utility in stratifying patients into distinct risk groups and providing prognostic insights that complement traditional RECIST assessments. Furthermore, we test the model’s robustness in data-sparse scenarios, specifically when cfDNA collection is limited to early treatment cycles. Finally, we address model interpretability by quantifying how individual molecular and clinical markers drive patient-specific predictions.

## Results

### A deep learning model for longitudinal ctDNA data

We present SHERLOC, a deep learning framework for survival analysis from longitudinal ctDNA data. SHERLOC integrates two complementary neural encoders applied to the IMpower150 cohort using serial liquid biopsy data at gene and panel levels (Figure 1a).

**Figure 1:**
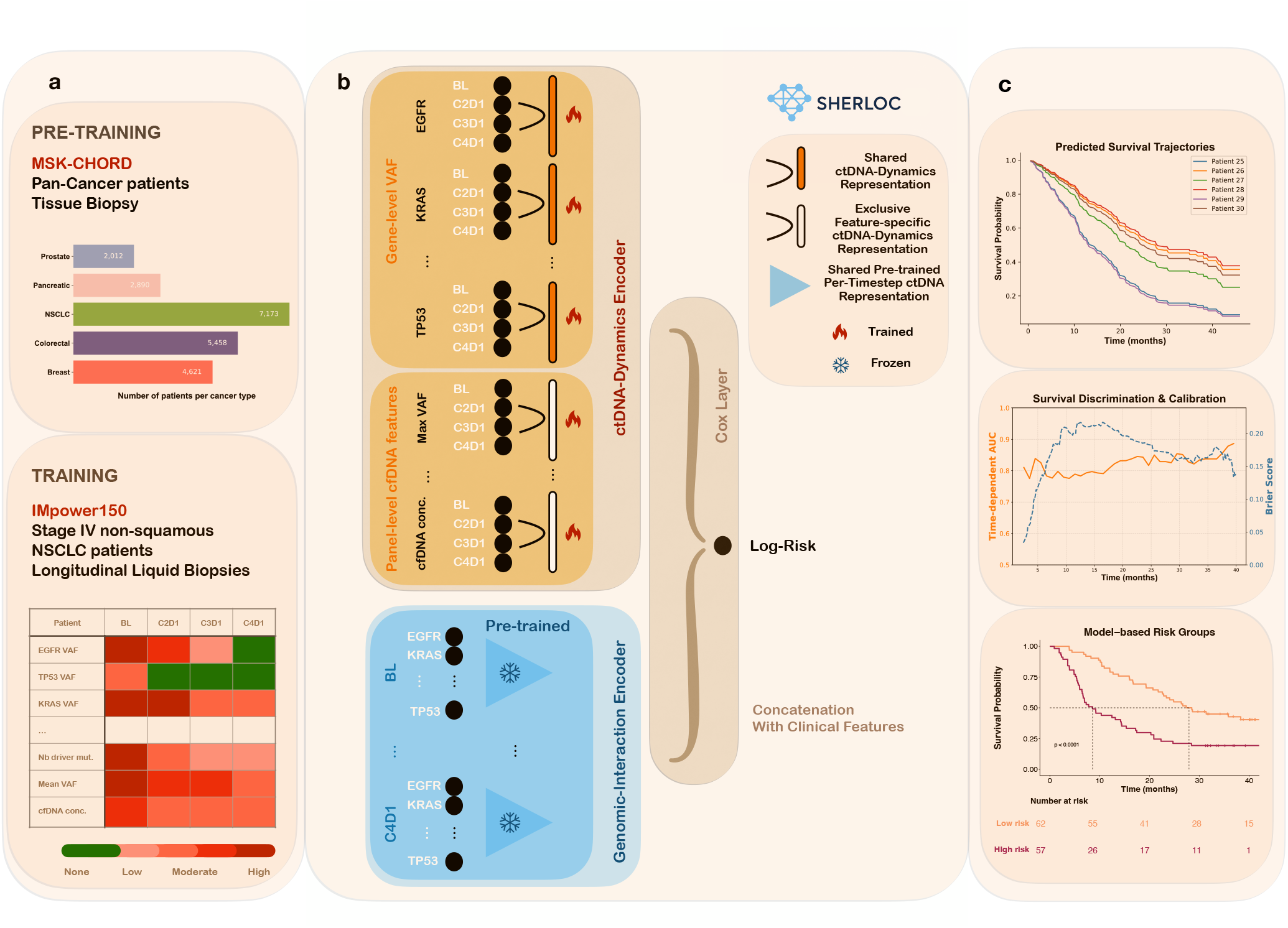
Overview of the SHERLOC framework. **(a) Data sources and pre-training.** A large-scale pan-cancer cohort (MSK-CHORD; *n* = 22, 068)—comprising tissue-biopsy variant allele frequencies (VAFs) and overall survival data—was utilized to pre-train a survival-aware, genomic-interaction encoder. The primary study data consists of 466 patients from the phase III IMpower150 non–small-cell lung cancer (NSCLC) trial, featuring up to four longitudinal ctDNA measurements per patient. **(b) Model architecture**. SHERLOC utilizes a dual-encoder strategy: (1) a ctDNA-dynamics encoder that learns shared temporal representations of gene-level VAFs and feature-specific temporal trajectories of panel-level biomarkers, and (2) a frozen, pre-trained genomic-interaction encoder designed to capture the prognostic interplay of gene alterations at fixed time points. The resulting representations from both encoders are concatenated with clinical variables and processed through an interpretable Cox proportional hazards framework. **(c) Clinical applications and task overview**. SHERLOC performs continuous-time survival analysis, including individual trajectory prediction, discrimination and calibration performance, and patient risk stratification to guide therapeutic interventions.

The *ctDNA-dynamics encoder* captures temporal patterns in variant allele frequencies (VAFs) across the first four treatment cycles (Figure 1b). A shared parameter strategy — where timestep-specific VAF coefficients are common across genes — reduces model complexity and enhances interpretability. The encoder also learns separate temporal representations for panel-level biomarkers (mean VAF, cfDNA concentration, maximum VAF, and number of driver mutations), accommodating their heterogeneous scales and biological interpretations.

The *genomic-interaction encoder* models cross-gene VAF interactions at fixed time points (Figure 1b). To overcome the limited sample sizes typical of liquid-biopsy trials, it is pre-trained on the large pan-cancer MSK-CHORD cohort [24] (22,068 patients, including 7,173 NSCLC) using a survival objective. Batch normalization mitigates systematic differences between ctDNA-derived and tissue-derived VAFs across datasets.

Representations from both encoders are concatenated with clinical features and passed to a Cox proportional hazards layer to generate individual risk scores. Together, SHERLOC enables survival discrimination, calibration, and patient stratification into interpretable risk groups to support therapeutic decision-making (Figure 1c).

An ablation study was performed to quantify the contribution of each encoder to survival discrimination and calibration. Model performance was assessed using the integrated Area under the ROC curve (iAUC) [26, 27], the Concordance index with inverse probability censoring weights (C-index) [28], and the integrated Brier score (IBS) [29]. To ensure the robustness and stability of the results, we adopted a repeated train–test split protocol with 100 repetitions (Monte Carlo cross-validation) [30].

As reported in Table 1, removing the ctDNA-dynamics encoder results in the most pronounced performance decline, with average iAUC dropping by 2.98, C-index by 2.95, and IBS increasing by 0.60, reflecting poorer discrimination and mean squared prediction error. This substantial drop highlights the dominant contribution of longitudinal ctDNA data to survival prediction and confirms that explicitly modeling treatment-induced temporal dynamics is critical for capturing prognostic signal.

**Table 1:** Ablation study results for SHERLOC on Overall Survival (OS). Performance is evaluated using time-to-event analysis with a repeated split train-test procedure (70% train / 30% test) over 100 random seeds; all metrics are reported on the held-out test sets. Values are reported as mean (SD), where SD denotes the corrected standard deviation as defined by the methodology of Nadeau and Bengio [25]. The integrated Area Under the Curve (iAUC) and Concordance index with inverse probability censoring weights (C-index) are presented on a 0–100 scale, with higher values indicating better survival discrimination. The Integrated Brier Score (IBS) is reported as IBS × 100, where lower values reflect a smaller mean squared prediction error.

| Model Variant | iAUC | C-index | IBS ($\times 100$) |
| --- | --- | --- | --- |
| SHERLOC (FULL) | **77.14** (2.28) | **69.64** (1.77) | **18.26** (0.52) |
| w/o ctDNA-dynamics encoder | 74.16 (2.84) | 66.69 (2.02) | 18.86 (0.53) |
| w/o genomic-interaction encoder | 74.89 (2.37) | 68.42 (1.76) | 18.55 (0.53) |
| w/o pre-training | 75.34 (2.38) | 68.55 (1.75) | 18.57 (0.51) |
| unfrozen genomic-interaction encoder | 75.83 (2.38) | 68.89 (1.79) | 18.45 (0.54) |
| w/o clinical features | 74.91 (2.56) | 67.89 (1.96) | 18.51 (0.58) |

Excluding the pre-trained genomic-interaction encoder also leads to a consistent decrease in performance (average iAUC *−*2.25, C-index *−*1.22, IBS +0.29), indicating that cross-gene interactions provide prognostic information complementary to ctDNA dynamics. A comparable drop is observed when the encoder is not pre-trained on the MSK-CHORD cohort (average iAUC *−*1.80, C-index *−*1.09, IBS +0.31), emphasizing the value of large-scale tumor DNA-derived data in improving generalization in small clinical cohorts. Moreover, unfreezing the pre-trained encoder during training on the IMpower150 cohort reduces performance (average iAUC *−*1.31, C-index *−*0.75), suggesting that treating the encoder as a frozen feature extractor effectively prevents representation drift and overfitting to the trial-specific data.

Finally, removing clinical covariates leads to a consistent yet moderate performance drop (a 2.23 decrease in iAUC and a 1.75 decrease in C-index, and a 0.25 increase in IBS). This indicates that while clinical features add complementary prognostic value, SHERLOC derives the majority of its predictive power from ctDNA-derived features.

Overall, the ablation analysis confirms that SHERLOC’s performance gains are driven primarily by its longitudinal modeling of ctDNA dynamics, with complementary prognostic contributions from cross-gene representations learned via pre-training on the MSK-CHORD cohort and from relevant clinical context.

### Comparative Performance Analysis: SHERLOC vs. Established Survival Frameworks

To rigorously assess the performance of the SHERLOC framework, we conducted a comprehensive comparative analysis against a broad spectrum of survival models, spanning traditional statistical methods to state-of-the-art deep learning models, as summarized in Table 2. Specifically, we compared SHERLOC against: (i) attention-based architectures [31], specifically the *Feature-wise* and *Temporal Transformer* models; (ii) recurrent neural networks, including *RNN* and *Gated Recurrent Unit (GRU)* [32] configurations; (iii) dense neural networks, such as *Multilayer Perceptron (DeepSurv)* [33]; (iv) traditional machine learning ensembles, such as *Random Survival Forest (RSF)* [34]; (v) regularized linear models, comprising Cox proportional hazards models with *Ridge* [35, 36], *Lasso* [37] and *Elastic Net* [38] penalization; (vi) hybrid methods, specifically *Functional Principal Component Random Forest (FPCRF)* [12], which utilizes functional PCA and random forest importance within an Elastic Net-penalized Cox framework; and (vii) Bayesian joint modeling via a multivariate *Joint Model* approach. To ensure a rigorous comparison, all benchmarked methods include the same frozen genomic-interaction encoder, with identical pre-training on the MSK-CHORD dataset. Detailed information regarding the methodology for each model is provided in the Methods section.

**Table 2:** Performance benchmark of SHERLOC against competing models. This table compares SHERLOC with various statistical, ensemble, and deep learning architectures for Overall Survival (OS). To ensure a controlled comparison, all methods integrated the same frozen genomic-interaction encoder, pre-trained on the pan-cancer MSK-CHORD cohort. Performance was evaluated using integrated Area Under the Curve (iAUC), Concordance index with inverse probability censoring weights (C-index), and Integrated Brier Score (IBS). All metrics were scaled to a 0–100 range for clarity; higher iAUC and C-index values indicate better survival discrimination, while lower IBS values reflect a smaller mean squared prediction error. For a rigorous measure of generalization, Monte Carlo cross-validation was performed. Results are reported as the mean performance across the 100 test sets obtained from random train–test splits, with corrected standard error calculated according to the Nadeau and Bengio methodology [25] to account for the overlap in training data during repeated splits.

| Method | iAUC | C-index | IBS |
| --- | --- | --- | --- |
| SHERLOC | **77.14 ± 2.28** | **69.64 ± 1.77** | **18.26 ± 0.52** |
| Feature-wise Transformer | 73.74 ± 3.01 | 66.49 ± 2.20 | 19.00 ± 0.56 |
| Temporal Transformer | 74.98 ± 2.49 | 67.49 ± 1.85 | 18.74 ± 0.54 |
| RNN | 74.07 ± 2.38 | 67.01 ± 1.77 | 18.88 ± 0.56 |
| GRU | 74.46 ± 2.51 | 67.27 ± 1.70 | 18.80 ± 0.52 |
| MLP (DeepSurv) | 74.58 ± 2.72 | 67.81 ± 2.02 | 18.73 ± 0.54 |
| RSF | 73.03 ± 2.38 | 66.35 ± 1.58 | 18.30 ± 0.67 |
| Ridge | 66.11 ± 3.01 | 61.59 ± 1.93 | 20.50 ± 0.46 |
| Lasso | 69.29 ± 3.79 | 63.82 ± 2.69 | 20.53 ± 3.56 |
| Elastic Net | 69.86 ± 3.18 | 64.28 ± 2.30 | 19.77 ± 0.74 |
| FPCRF | 73.00 ± 3.05 | 66.26 ± 2.18 | 18.66 ± 0.83 |
| Joint Model | 72.13 ± 5.77 | — | 20.67 ± 2.48 |

SHERLOC achieves the best performance across all three evaluation metrics, with an iAUC of 77.14 ± 2.28, a C-index of 69.64 ± 1.77, and an IBS of 18.26 ± 0.52, consistently outperforming both deep learning–based and classical approaches. Within the neural network models, Transformer-based and DeepSurv architectures achieve reasonably competitive discrimination (iAUC ≈73.7– 75.0; C-index ≈66.5–67.8) and mean squared prediction error (IBS ≈18.7–19.0). Multilayer Perceptron (DeepSurv) employs feature-specific representations for each longitudinal cfDNA/ctDNA feature, while Transformer-based models use shared projection matrices to generate queries, keys, and values for each token. In contrast, SHER-LOC adopts a hybrid strategy, learning a shared representation for gene-level VAFs alongside feature-specific representations for panel-level cfDNA and ctDNA features, thereby achieving a more effective balance between shared and specialized modeling.

Recurrent models (RNN and GRU) show comparable performance to Transformer-based architectures, with iAUC values between 74.1 and 74.5, C-index values around 67.0–67.3, and IBS values around 18.8–18.9. This overlap in performance is plausibly ex-plained by the limited temporal depth of the ctDNA data, which comprises only four longitudinal time points. In such short sequences, the advantages of recurrence for modeling long-range temporal dependencies are constrained, limiting potential gains in both discrimination and calibration.

Random Survival Forest (RSF) demonstrates moderate discriminative performance (iAUC = 73.03 ± 2.38; C-index = 66.35 ± 1.58). This limitation may stem from the reduced ability of fixed-tree ensemble methods to effectively capture temporally structured dynamics in longitudinal data. Nevertheless, RSF exhibits competitive mean squared prediction error, as indicated by an IBS of 18.30 *±* 0.67.

Linear Cox proportional hazards models with conventional penalization (Ridge, Lasso, and Elastic Net) exhibit substantially lower discriminative performance (iAUC ≈66.1–69.9; C-index ≈61.6– 64.3) and higher IBS values (≈19.8–20.5), indicating less accurate risk ranking and a larger mean squared distance between predicted survival probabilities and actual observed outcomes. This reduced performance is likely attributable to their limited ability to model non-linear relationships and complex feature dependencies inherent in longitudinal ctDNA data.

*Functional principal component random forest (FPCRF)* combines functional principal component extraction with random forest– based variable selection, followed by dimension reduction and an Elastic Net–regularized Cox model [12]. In terms of performance, FPCRF demonstrates moderate survival discrimination (iAUC 73.00 *±* 3.05, C-index 66.26 *±* 2.18), along with reasonable mean squared prediction error (IBS 18.66 *±* 0.83).

The Bayesian multivariate *Joint Model* approach shows average performance in survival discrimination (iAUC = 72.13 ± 5.77); its high IBS (20.67 ± 2.48) highlights the inherent difficulty of using joint models in sparse, high-dimensional longitudinal settings.

### SHERLOC ctDNA-based risk score improves upon standard RECIST criteria

To evaluate the independent prognostic value of the SHERLOC ctDNA-based risk score, we performed a multivariable Cox proportional hazards analysis (Figure 2a; Supplementary Figure 1b). In the test set (*N* = 119), the SHERLOC ctDNA risk score—derived only from longitudinal ctDNA data collected at BL, C2D1, and C3D1—emerged as a highly significant independent predictor of overall survival, even when adjusted for established clinical features and C3D1 radiographic assessments via RECIST v1.1 [1]. When treated as a continuous variable, the score yielded a Hazard Ratio (HR) of 3.60 (95% confidence interval [CI]: 1.75–7.41; *p <* 0.001), independent of whether a patient was classified as having Partial Response (PR), Stable Disease (SD) or Progressive Disease (PD) by conventional imaging. While RECIST categories were prognostic—with PD showing an HR of 3.82 (95% CI: 1.25-11.67; *p* = 0.02) and SD showing an HR of 1.82 (95% CI: 1.06-3.13; *p* = 0.03) compared to the Partial Response (PR) reference—the SHERLOC score provided independent additive value, suggesting that molecular changes in ctDNA capture prognostic disease risk that is not yet visible on a CT scan. Among clinical variables, ECOG status remained a strong clinical predictor (HR: 3.17, 95% CI: 1.92-5.24 ; *p <* 0.001), while female sex was associated with improved survival (HR: 0.58, 95% CI: 0.35-0.95; *p* = 0.03).

**Figure 2:**
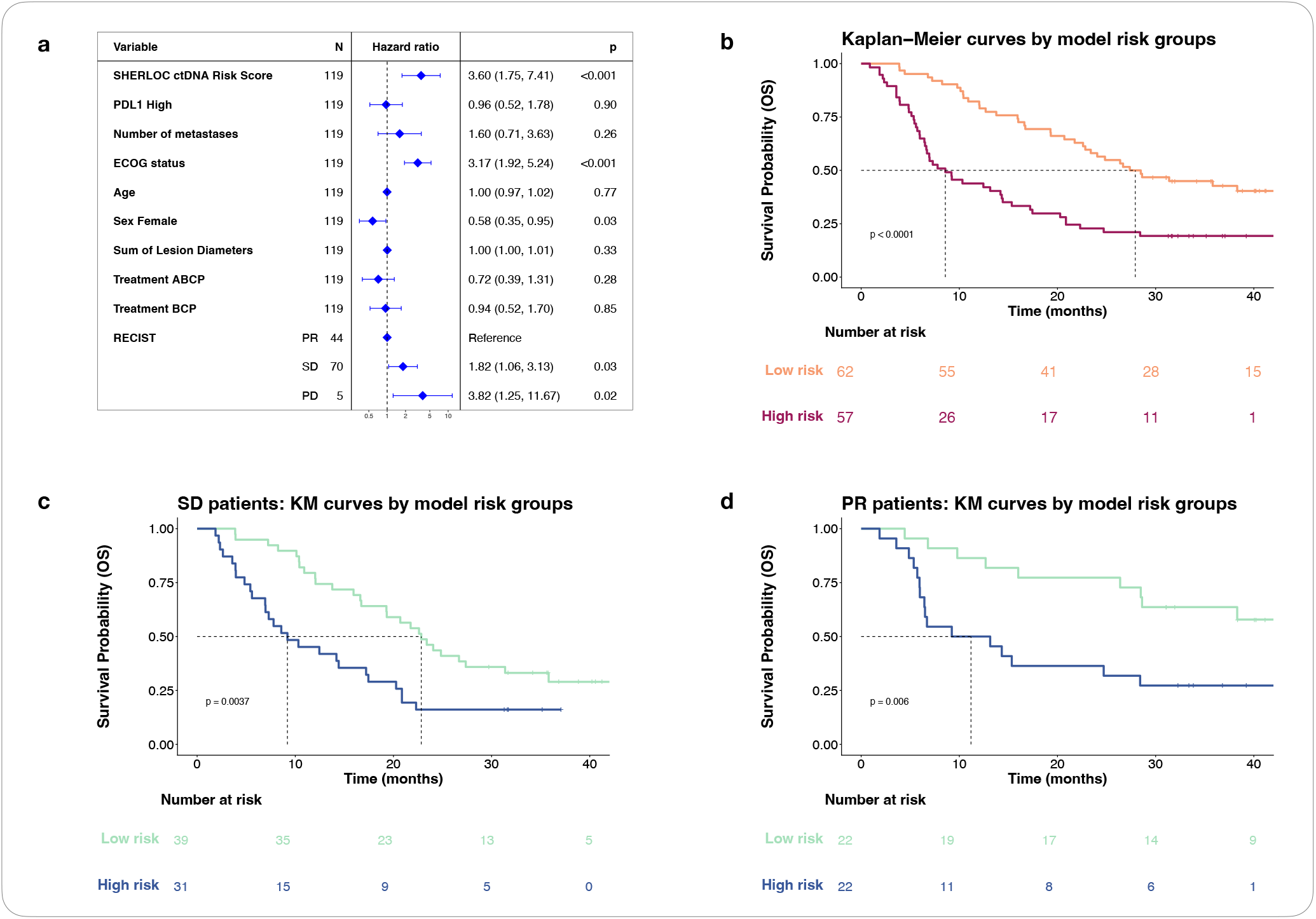
SHERLOC ctDNA-based risk model improves upon traditional RECIST criteria. **(a)** Overall Survival (OS) hazard ratio (HR) of a multivariable Cox regression in the test set (*n* = 119) incorporating clinical features, RECIST categories assessed at C3D1 and ctDNA-based model risk score derived from SHERLOC trained only on ctDNA samples collected from baseline to C3D1. OS was recalculated from the RECIST assessment date (landmark analysis). Of 398 patients remaining after pre-landmark OS exclusion, 395 had C3D1 RECIST assessments. **(b)** OS Kaplan-Meier curves in the test set stratified by the SHERLOC ctDNA-based risk categories. Median risk scores of the training set (*n* = 276) were used as a cut point to derive low- and high-risk patients. **(c, d)** OS Kaplan-Meier curves in the test set for patients categorized as SD **(c)** or PR **(d)** from RECIST and stratified by the ctDNA-based model risk categories. **Abbreviations:** CR, complete response; PD, progressive disease; PR, partial response; SD, stable disease.

To facilitate clinical interpretability, a binary risk threshold was defined using the median SHERLOC risk score derived from the training cohort (*N* = 279). Patients were subsequently classified into Low Risk (below the threshold) and High Risk (above the threshold) groups (Supplementary Figures 1a, 1c). In the test set, patients classified as High Risk (*n* = 57) exhibited a significantly increased hazard of death compared with those in the Low Risk reference group (*n* = 62). Specifically, High Risk patients had more than twice the risk of death (HR = 2.18, 95% CI: 1.33–3.57; *p* = 0.002), corresponding to a 118% increase in risk, after adjustment for relevant clinical covariates including ECOG performance status and RECIST radiographic response (Supplementary Figure 1a). In the test cohort, the median OS for patients classified as Low Risk was 27.9 months, compared with 8.6 months for those classified as High Risk (*p <* 0.0001; Figure 2b), demonstrating robust prognostic discrimination. Within patients with uniform RECIST response, the SHER-LOC ctDNA-based score identified risk groups with distinct survival (Figures 2c, 2d; Supplementary Figures 1e, 1f). Among patients with stable disease (SD), median OS was 22.8 months for the Low Risk group, whereas High Risk patients had a median OS of 9.2 months (*p* = 0.0037; Figure 2c). Similarly, among with partial response (PR), median OS was not reached in the Low Risk group, and 11.2 months in the High Risk group (*p* = 0.006; Figure 2d).

Collectively, these findings demonstrate that the SHERLOC ctDNA-based risk score enables consistent and clinically meaningful stratification of patient outcomes across the overall cohort and within RECIST-defined subgroups, supporting its potential utility as a prognostic tool to inform treatment decision-making.

### Model robustness to reduced longitudinal cfDNA samples

In this section, we evaluate the performance of survival models when the number of longitudinal cfDNA measurements per patient is reduced from four to three (BL, C2D1, C3D1) or two time points (BL, C2D1), as reported in Table 3. This setting reflects real-world clinical constraints, where the substantial costs associated with NGS-based profiling of cfDNA often prohibit frequent, high-density monitoring. Using the repeated train–test split protocol with 100 repetitions (Monte Carlo cross-validation) [30], the benchmark reports the mean iAUC, C-index, and IBS across the 100 test sets, together with corrected standard deviations following the methodology of Nadeau and Bengio [25].

**Table 3:**
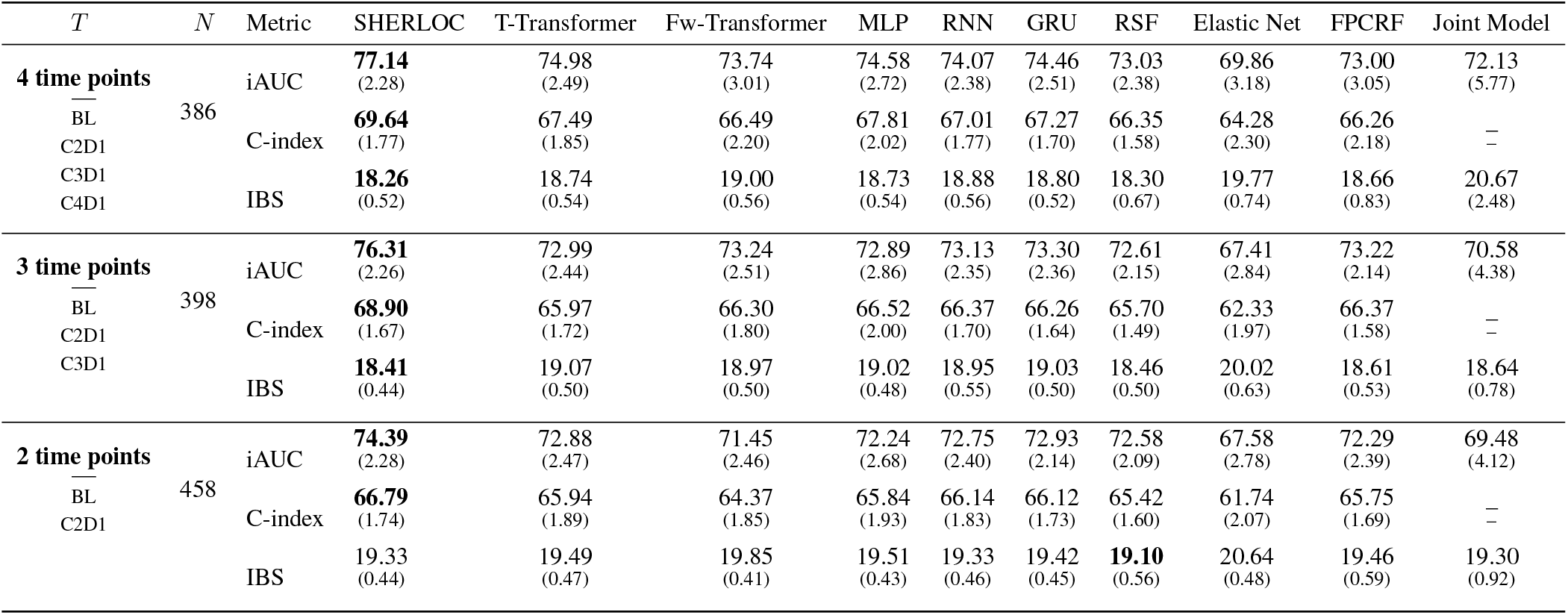
Robustness to reduced longitudinal ctDNA measurements. Performance comparison of diverse statistical, ensemble, and deep learning models against SHERLOC under constrained temporal data for Overall Survival (OS). Evaluation focuses on scenarios where ctDNA collection was restricted to the first two, three, or four treatment cycles of treatment (*T* = 2, 3, 4). To ensure parity, all models utilized a common frozen genomic-interaction encoder, pre-trained on the MSK-CHORD cohort. Metrics include integrated Area Under the Curve (iAUC), Concordance index with inverse probability censoring weights (C-index), and Integrated Brier Score (IBS). All metrics were scaled to a 0–100 range for clarity. For a robust assessment of generalization, we utilized Monte Carlo cross-validation. Reported values represent the mean performance across the 100 test sets obtained from random train–test splits; along with the corrected standard error derived using the Nadeau and Bengio method.

When three longitudinal cfDNA measurements per patient are available (*N* =398 patients), SHERLOC maintains the best overall performance, achieving an iAUC of 76.31 ±2.26, a C-index of 68.90 ± 1.67, and an IBS of 18.41 ± 0.44, indicating strong discriminative ability and well-calibrated survival predictions. Among the other neural architectures, GRU and RNN demonstrate notable robustness, with an iAUC of 73.30 ± 2.36 and 73.13 ± 2.35 respectively, and a C-index of 66.26 ± 1.64 and 66.37 ± 1.70. The Feature-wise Transformer, which applies attention across longitudinal cfDNA/ctDNA feature tokens, achieves closely competitive performance (iAUC = 73.24 *±* 2.51; C-index = 66.30 *±* 1.80; IBS = 18.97 *±* 0.50). The Temporal Transformer, which instead applies attention over timestep tokens, shows slightly lower discrimination (iAUC = 72.99 ± 2.44; C-index = 65.97 ± 1.72; IBS = 19.07 ±0.50), and DeepSurv yields comparable figures (iAUC = 72.89 ± 2.86; C-index = 66.52 ± 2.00; IBS = 19.02 ± 0.48). IBS values remain broadly consistent across the competing neural network models (≈ 19.0 ± 0.5), in contrast to SHERLOC (18.41 ± 0.44), which reflects a lower mean squared prediction error. Random Survival Forest exhibits more moderate discriminative performance compared with neural network survival models (iAUC = 72. ± 61 2.15, C-index = 65.70 ± 1.49), but maintains relatively favorable IBS values (IBS = 18.46 ± 0.50), suggesting that while RSF is less effective at risk ranking, its mean squared prediction error remains comparatively robust. The Bayesian multivariate joint model demonstrates inferior survival discrimination (iAUC = 70.58 ± 4.38) but remains competitive in terms of IBS (18.64 ± 0.78). Linear Cox proportional hazards models with Elastic Net penalization demonstrate markedly lower discrimination and consistently higher IBS values. FPCRF achieves improved survival discrimination and mean squared prediction error compared with a standard Elastic Net Cox model (iAUC: 73.22 ± 2.14 vs. 67.41 ± 2.84, C-index: 66.37 ± 1.58 vs. 62.33 ± 1.97, IBS 18.61 ± 0.53 vs. 20.02 ± 0.63), which can be attributed to random forest–based importance feature selection prior to integration into the Elastic Net Cox model, while capturing longitudinal temporal patterns through functional principal component representations.

Restricting the analysis to two longitudinal cfDNA measurements per patient (*N* = 458) causes a systematic drop in performance across all models. This degradation highlights the difficulty of capturing complex, patient-specific temporal dynamics from just two early-stage timepoints. In this constrained setting, SHERLOC nevertheless maintains the strongest overall performance profile, demonstrating superior discriminative accuracy (iAUC = 74.39 ± 2.28, C-index = 66.79 ± 1.74) while maintaining a competitive mean squared prediction error (IBS = 19.33 ± 0.44), confirming its robustness to sparse longitudinal inputs. Under these conditions, GRU, RNN, and the Temporal Transformer achieve competitive performance across both survival discrimination and calibration-related aspects (iAUC ≈ 72.8–72.9; C-index ≈ 65.9–66.1; IBS ≈ 19.3–19.5), indicating relative robustness even with only two time points. By contrast, the Feature-wise Transformer (iAUC = 71.45 ± 2.46; C-index = 64.37 ± 1.85; IBS = 19.85 ± 0.41) and DeepSurv (iAUC = 72.24 ± 2.68; C-index = 65.84 ± 1.93; IBS = 19.51 ± 0.43) show more pronounced degradation relative to the three- and four-timepoint settings. Random Survival Forest achieves relatively robust discriminative ability (iAUC = 72.58 ± 2.09; C-index =65.42 ± 1.60) and records the lowest overall prediction error, with an IBS of 19.10 ± 0.56. This is consistent with the intuition that RSF’s primary limitation lies in capturing temporal dynamics — a constraint that becomes less penalizing when longitudinal depth is drastically reduced to two time points — while its tree-based probability estimation remains well-calibrated regardless. The Bayesian multivariate joint model yields lower discriminative ability (iAUC = 69.48 ± 4.12) but maintains a competitive IBS (19.30 ± 0.92). Finally, Elastic Net (iAUC = 67.58 ± 2.78; C-index = 61.74 ± 2.07; IBS = 20.64 ± 0.48) continues to show consistently low performance, as it does across all timepoint settings, reflecting its structural inability to capture non-linear temporal dynamics rather than a specific sensitivity to data sparsity. FPCRF (iAUC = 72.29 ± 2.39; C-index = 65.75 ± 1.69; IBS = 19.46 ± 0.59) retains reasonable discrimination, further illustrating that some capacity to model functional structure remains beneficial even with minimal longitudinal data. Overall, the performance gaps between SHERLOC and competing methods tend to narrow as the longitudinal information is progressively constrained, particularly when only two longitudinal cfDNA measurements per patient are available. This trend is expected, as reducing the temporal depth limits the amount of dynamic information that can be exploited by more expressive temporal models. Notably, all methods share the same static genomic-interaction encoder with identical pre-training, ensuring a fair comparison. Performance differences therefore primarily reflect the models’ ability to exploit longitudinal structure, which is inherently limited with two time points, leading to more similar performance across methods.

### Interpretability of representations in SHERLOC

The architecture of the SHERLOC model enables granular decomposition of prognostic signals by fusing representations from the ctDNA-dynamics encoder, the genomic-interaction encoder, and clinical features into a single fully connected Cox proportional hazards log-risk layer. To evaluate the individual contribution of each feature, we implemented a robust validation framework where learned representations were frozen and the final fully connected layer was re-trained using a resampling procedure. Statistical significance was rigorously determined via the corrected *t*-test and standard deviation, following the methodology of Nadeau and Bengio [25], to account for the lack of independence inherent in overlapping training sets.

Analysis of longitudinal panel-level biomarkers revealed that SHERLOC identifies several cfDNA-derived features as significant predictors of overall survival (OS), as reported in Figure 3a. By learning feature-specific temporal representations, the model weighs clinical importance across treatment cycles (BL, C2D1, C3D1, and C4D1). Specifically, longitudinal Mean VAF representations were significantly associated with poorer survival ([1] : *β* = 0.289, *HR* = 1.34 95% CI [1.17, 1.52], *p <* 0.001; [2] : *β* = 0.373, *HR* = 1.45 [1.25, 1.69], *p <* 0.001), with weightings concentrated at the C3D1 time point. This suggests that tumor burden after two therapy cycles is a critical prognostic indicator. Similarly, longitudinal cfDNA concentration representations showed a strong positive association with mortality risk ([1] : *β* = 0.430, *HR* = 1.54 [1.31, 1.80], *p <* 0.001; [2] : *β* = 0.396, *HR* = 1.49 [1.27, 1.74], *p <* 0.001), primarily weighted at C4D1. Notably, both longitudinal representations of the number of driver mutations exhibited a consistent temporal shift, transitioning from negative weighting at baseline to strong positive weighting at C4D1. These results indicate that an increase in the driver mutation count during therapy is significantly associated with poorer survival ([1] : *β* = 0.52, *HR* = 1.68 [1.47, 1.93], *p <* 0.001; [2] : *β* = 0.52, *HR* = 1.68 [1.46, 1.94], *p <* 0.001).

**Figure 3:**
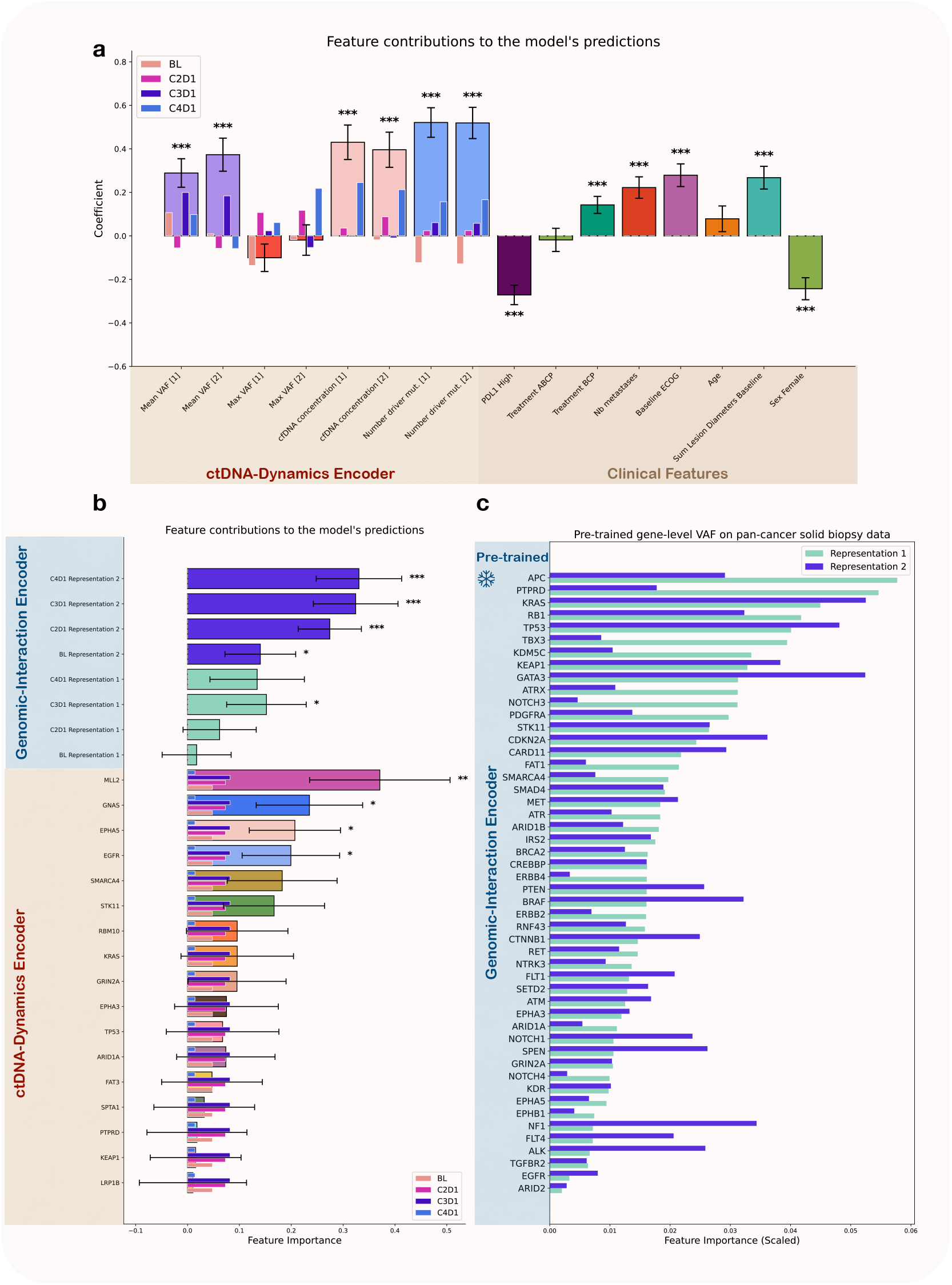
Interpretation of SHERLOC representations for Overall Survival (OS). **(a) Panel-level cfDNA-derived representations and clinical features.** Bar plots display the predictive coefficients of the SHERLOC output Cox layer for two longitudinal representations per panel-level cfDNA feature (e.g., Mean VAF [1] and [2]) as well as for clinical variables. Empirical distributions of coefficients were derived from a resampling-based validation framework. Statistical significance was assessed using a corrected resampled *t*-test [25] (* *p <* 0.05, ** *p <* 0.01, *** *p <* 0.001). The inset colored bars indicate the weighting of specific timesteps within the longitudinal representations. **(b) Gene-level VAFs and genomic-interaction representations**. Bar plots display the coefficients for gene-level VAFs and pre-trained per-timestep ctDNA representations. Mean coefficients were reported in absolute values (i.e. feature importance) to avoid misinterpretation (see Methods), while statistical significance was assessed on raw coefficients using a corrected resampled *t*-test (* *p <* 0.05, ** *p <* 0.01, *** *p <* 0.001). **(c) Gene contributions in the genomic-interaction encoder**. Bar plots indicate the percentage contribution of each gene to its respective representation in the genomic-interaction encoder. These two representations of gene-level VAFs were learned in the pre-training phase on the MSK-CHORD pan-cancer cohort, restricted to the genes shared with the IMpower150 trial sequencing panel.

Regarding clinical covariates, consistent with the original trial analysis [10], high PD-L1 expression was significantly associated with improved survival (*β* = *−*0.272, *HR* = 0.76 [0.70–0.83], *p <* 0.001). The BCP regimen was associated with significantly poorer survival outcomes compared to the ACP arm (*β* = 0.142, *HR* = 1.15 [1.07–1.25], *p <* 0.001), whereas the ABCP arm showed a non-significant trend toward improved survival relative to the ACP group (*β* = *−*0.018, *HR* = 0.98 [0.88–1.09], *p* = 0.730). Although ABCP and ACP arms demonstrated similar overall survival in the original analysis [10], the U.S. FDA approval was specifically based on the superiority of ABCP over BCP [15, 39], in line with our findings. Demographic and disease burden factors also aligned with prior results [10]: female sex was associated with better survival (*β* = *−*0.243, *HR* = 0.78 [0.71–0.87], *p <* 0.001), while increased sum of lesion diameters (*β* = 0.268, *HR* = 1.31 [1.18–1.45], *p <* 0.001), higher metastasis count (*β* = 0.222, *HR* = 1.25 [1.13–1.38], *p <* 0.001), and elevated Baseline ECOG scores (*β* = 0.279, *HR* = 1.32 [1.19–1.47], *p <* 0.001) were all predictors of poorer survival.

Regarding gene-specific longitudinal dynamics, the ctDNA-dynamics encoder applies a shared temporal mask across individual gene VAFs, prioritizing the C2D1 and C3D1 time points. To evaluate the contribution of gene-level VAFs across our architecture, we analyze feature importance (see Methods) within both the longitudinal ctDNA-dynamics encoder and the pre-trained genomic-interaction encoder, as illustrated in Figures 3b and 3c. Several gene representations demonstrated independent prognostic value (Figure 3b). The longitudinal representation of *MLL*2 (*KMT* 2*D*) showed significant association with survival (*p* = 0.007), as well as the representation of *GNAS* (*p* = 0.024), *EPHA*5 (*p* = 0.021), and *EGFR* (*p* = 0.037). Finally, SHERLOC leverages the MSK-CHORD pancancer cohort to pre-train a genomic-interaction encoder capturing cross-gene prognostic interactions. Two latent representations were extracted with distinct genomic profiles. Representation 1, primarily driven by *APC, PTPRD*, and *KRAS* (Figure 3c), reached statistical significance specifically at C3D1 (*p* = 0.049). In contrast, Representation 2, heavily influenced by the *KRAS/TP* 53*/GATA*3*/ KEAP* 1 axis (Figure 3c), demonstrated a cumulative prognostic effect. Its association with OS increased from Baseline (*p* = 0.042) to C4D1 (*p <* 0.001).

## Discussion

In this study, we addressed a central methodological and clinical challenge in contemporary liquid biopsy research: how to extract robust, interpretable, and clinically actionable prognostic information from high-dimensional ctDNA measurements collected at only a few longitudinal time points within the constraints of a clinical trial. Using cfDNA data from IMpower150, we introduced SHERLOC, a deep learning framework specifically designed for short longitudinal trajectories, limited sample sizes, and strong interpretability requirements. Our results demonstrate that SHERLOC consistently outperforms a broad range of classical statistical methods, machine learning approaches, and state-of-the-art deep learning models in survival discrimination and calibration, while remaining robust to reductions in the number of available liquid biopsy time points. Importantly, SHERLOC adopts a hybrid longitudinal modeling strategy: it learns a shared temporal structure across gene-level VAFs, while simultaneously learning feature-specific temporal representations for panel-level cfDNA/ctDNA biomarkers. This design enables an effective balance between shared and specialized modeling, substantially reducing model complexity for high-dimensional gene-level data while preserving flexibility to capture heterogeneous longitudinal behaviour across biologically distinct panel-level cfDNA-derived features.

A key finding of this work is the dominant prognostic value of longitudinal ctDNA data under therapy. The ablation analysis showed that removing the ctDNA-dynamics encoder resulted in the largest degradation in model performance, underscoring that temporal changes in ctDNA – rather than static molecular snapshots alone – encode critical information about treatment response and disease evolution. Beyond temporal dynamics, our results highlight the value of integrating external large-scale genomic resources through pre-training. The genomic-interaction encoder, pre-trained on the pan-cancer MSK-CHORD cohort, provided complementary prognostic signal beyond trial-specific data. These findings suggest that survival-aware representations learned from large tissue-based sequencing cohorts can be successfully transferred to ctDNA-based analyses, despite differences in assay technology and biological context, provided that appropriate batch normalization strategies are applied. More broadly, this approach offers a principled way to maximize existing resources by leveraging widely available solid-tissue cohorts to bridge the gap created by the smaller sample sizes of more recent liquid-biopsy clinical trials.

The demonstrated prognostic value of the SHERLOC ctDNA-based risk score underscores its potential clinical utility as an early, non-invasive tool for patient stratification. Importantly, despite being derived solely from ctDNA measurements, the risk score remained a strong and independent predictor of overall survival even after adjustment for established clinical covariates and standard radiographic response. This finding highlights the ability of molecular dynamics captured in plasma to reflect evolving disease biology that may precede or complement imaging-based assessments. The added value of SHERLOC within homogeneous RECIST response groups further illustrates its capacity to resolve clinically meaningful heterogeneity among patients who would otherwise be classified as having similar treatment responses. Such early identification of high-risk patients despite apparent radiographic stability or response could enable more timely treatment adaptation, intensified monitoring, or enrollment into alternative therapeutic strategies.

A central strength of SHERLOC is its ability to provide clinically and biologically interpretable insights alongside strong predictive performance. By constraining the final prediction to a single Cox proportional hazards layer and rigorously validating feature contributions through repeated resampling, the model enables robust attribution of prognostic signal across gene- and panel-level longitudinal cfDNA data, and clinical covariates. The interpretability analysis highlights that feature-specific temporal representations of panel-level cfDNA biomarkers—particularly mean VAF at C3D1, cfDNA concentration at C4D1, and the evolving number of driver mutations between baseline and C4D1—are consistently the dominant contributors to survival prediction. Furthermore, longitudinal VAF representations of specific genes (including *KMT* 2*D* [*MLL*2], *GNAS, EPHA*5 and *EGFR*) emerges as drivers of the model’s output. Representation from the genomic-interaction encoder, pre-trained on the MSK-CHORD cohort and primarily driven by the *KRAS/ TP* 53*/GATA*3*/KEAP* 1 axis, exhibits a cumulative prognostic effect that intensifies as therapy progresses. Importantly, SHER-LOC recapitulates established clinical associations observed in IMpower150 original clinical trial analysis [10]. Together, these results demonstrate that SHERLOC achieves interpretability not through post hoc explanation, but through architectural design, yielding a transparent and statistically grounded framework that links longitudinal ctDNA evolution to patient prognosis in a manner that is both clinically credible and methodologically robust.

Several limitations merit consideration. First, this analysis was conducted within a single randomized clinical trial in advanced non-squamous NSCLC, and external validation in independent cohorts will be essential to assess generalizability to other cancer types. Second, while SHERLOC was designed to accommodate short longitudinal sequences of ctDNA data, it does not explicitly model irregular sampling, which may be relevant in broader real-world settings. Finally, although this study emphasizes interpretability at the level of learned longitudinal representations of gene- and panel-level ctDNA features, further work is needed to translate these insights into simple clinical rules or decision-support tools.

In conclusion, this work shows that meaningful and interpretable survival modeling can effectively address the unique challenges of on-treatment longitudinal ctDNA analysis in clinical trials with limited sample sizes by explicitly tailoring the deep learning architecture to the statistical structure of the data. Across extensive benchmarking, SHERLOC consistently demonstrated superior discrimination, calibration, and robustness compared with both classical statistical approaches and state-of-the-art deep learning models, while maintaining interpretability through a transparent survival modeling framework. Importantly, the resulting ctDNA-based risk score provides prognostic information that complements standard imaging-based response assessments, underscoring the clinical potential of longitudinal ctDNA dynamics as an early and sensitive indicator of treatment benefit.

## Methods

### Study Population and Landmark Analysis

Patient data were derived from the Phase 3 IMpower150 trial (NCT02366143), which randomized chemotherapy-naive patients with metastatic nonsquamous NSCLC to receive every 3 weeks atezolizumab plus carboplatin and paclitaxel with (ABCP) or without (ACP) bevacizumab, or bevacizumab plus chemotherapy (BCP).1 To evaluate the association between longitudinal ctDNA dynamics and clinical outcomes, we employed a landmark analysis (excluding joint models) to mitigate immortal time bias. Overall survival (OS) was calculated from the time of the final on-treatment ctDNA collection, with patients experiencing an event prior to the landmark excluded. The cohort evaluating plasma dynamics from baseline (BL) to Cycle 2 Day 1 (C2D1) included 458 patients, while the cohorts extending to C3D1 and C4D1 comprised 398 and 386 patients, respectively.

### ctDNA Sequencing and Variant Filtering

Cell-free DNA (cfDNA) was longitudinally collected from 466 patients in the IMpower150 study. For this analysis, evaluable patients (e.g., *n* = 386 for the C4D1 landmark) were required to have available plasma at baseline and all subsequent on-treatment time points (C2D1, C3D1, and C4D1), along with matched peripheral blood mononuclear cells (PBMCs). Baseline samples were sequenced using a prototype FoundationOne Liquid CDx assay (FMI), covering 394 genes (*>* 1.25 Mb). To enhance sensitivity and cost-efficiency for longitudinal tracking, on-treatment samples were analyzed using a high-depth custom fixed-panel assay. To ensure high-fidelity tumor variant calling, matched PBMCs were utilized to algorithmically filter rare germline variants and clonal hematopoiesis of indeterminate potential (CHIP), including mutations in canonical driver genes [10].

### MSK-CHORD Cohort and Pre-processing

We leveraged the pan-cancer MSK-CHORD cohort [24] for model pre-training. Gene-level variant allele frequencies (VAFs) were extracted for 59 genes overlapping with those sequenced in the IMpower150 cohort to ensure cross-dataset compatibility. Silent mutations were excluded, and genes mutated in fewer than 2% of patients were removed to reduce sparsity and limit the influence of rare events.

After applying these filters, the final MSK-CHORD population comprised 22,068 patients, including 7,173 with non-small cell lung cancer, 5,458 with colorectal cancer, 4,621 with breast cancer, 2,890 with pancreatic cancer, and 2,012 with prostate cancer. Overall survival data were available for all individuals and were used as endpoints during the pre-training phase to learn survival-aware genomic representations.

### SHERLOC Architecture and Development

The SHERLOC deep learning framework was designed as an end-to-end joint neural network to integrate longitudinal liquid biopsy data with baseline clinical characteristics for survival analysis. The architecture comprises two primary encoding branches: a ctDNA-dynamics encoder and a pre-trained genomic-interaction encoder. Throughout the network, the sigmoid activation function, defined as *σ*(*z*) = (1 + *e*^*−z*^)^*−*1^, is applied to introduce non-linearity.

#### ctDNA-Dynamics Encoder

The ctDNA-dynamics encoder processes temporal genomic data across the first four cycles of therapy: baseline (*BL*), *C*2*D*1, *C*3*D*1, and *C*4*D*1. For gene-level inputs, we included variant allele frequencies (VAFs) for 17 genes mutated in at least 5% of the cohort at baseline (*ARID1A, EGFR, EPHA3, EPHA5, FAT3, GNAS, GRIN2A, KEAP1, KRAS, LRP1B, MLL2, PTPRD, RBM10, SMARCA4, SPTA1, STK11*, and *TP53*). These inputs undergo batch normalization followed by a one-dimensional temporal convolution. This operation reduces the sequence of four time points *x*_*g*_ = [*x*_*BL*_, *x*_*C*2_, *x*_*C*3_, *x*_*C*4_] into a single shared representation *y*_*g*_ via the weighted sum:

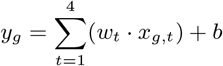

where *w* and *b* represent learnable temporal kernels and bias parameters shared across genes. Activation functions applied are sigmoid activations.

Simultaneously, four panel-level features—mean VAF, maximum VAF, cfDNA concentration, and the number of detected driver mutations—are processed. Each feature *x*_*p*_ ∈ ℝ^4^ is mapped to a unique two-dimensional representation *z*_*p*_ *∈* ℝ^2^ using a feature-specific linear transformation:

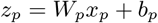

where *W*_*p*_ ∈ ℝ^2*×*4^ is the weight matrix specific to each panel feature.

#### Pre-trained genomic-interaction encoder

To leverage larger external datasets, we implemented a genomic-interaction encoder pre-trained on solid biopsy VAF data from 59 genes common to both the study cohort and the MSK-CHORD cohort. This encoder utilizes a linear transformation to map the gene vector *v*_*t*_ ∈ ℝ^59^ at a given timestep *t* to a two-dimensional embedding *e*_*t*_:

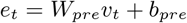

where *W*_*pre*_ ∈ ℝ^2*×*59^ and *b*_*pre*_ are frozen parameters. This transformation is applied identically across each individual timestep (*BL, C*2*D*1, *C*3*D*1, *C*4*D*1), resulting in a total of eight output features from this branch. Activation functions applied are sigmoid activations.

The final stage of the network involves the concatenation of the dynamic representations (*y*_*g*_, *z*_*p*_), the frozen embeddings (*e*_*t*_), and baseline clinical variables *c* (including PD-L1 expression, ECOG status, treatment arm, and number of metastatic sites). Prior to concatenation, features undergo batch normalization for numerical stability. A drop-out layer was then applied with probability equal to 0.4. The resulting unified feature vector Φ is passed to a final linear layer to calculate the predicted log-risk *η*:

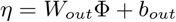

The model was trained by minimizing the negative log-partial likelihood of the Cox proportional hazards model:

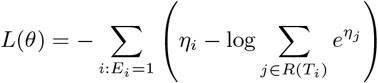

where *E*_*i*_ is the event indicator and *R*(*T*_*i*_) is the set of patients at risk at time *T*_*i*_.

### Transformer Architectures

To explore the impact of feature-wise versus temporal-wise attention, we implemented two distinct Transformer-based configurations: *Temporal Transformer* and *Feature-wise Transformer*. The input data is structured as a time-series 𝒳∈ ℝ^*T ×C*^, where *T* = 4 represents the clinical timesteps and *C* = 21 represents the combined longitudinal channels (17 gene-level VAFs and 4 panel-level features).

#### Temporal Transformer

The *Temporal Transformer* configuration utilizes the input data, 𝒳∈ ℝ^*T ×C*^, treating each of the four clinical time points as a single token containing *C* features. This setup prioritizes the temporal evolution of the liquid biopsy profile as a whole. The self-attention mechanism computes dependencies between different stages of therapy (e.g., the relationship between baseline profiles and *C*4*D*1 responses), effectively learning a non-linear weighting of the treatment timeline. The output of the temporal Transformer is fused with the frozen genomic-interaction encoder outputs and baseline clinical characteristics *c* to compute the final unified feature vector Φ for Cox proportional hazards regression. Prior to concatenation, features undergo batch normalization to ensure numerical stability, followed by a dropout layer with a regularizing probability of 0.4—consistent with all competing neural network frameworks.

#### Feature-wise Transformer

In the *Feature-wise Transformer* configuration, the model treats each individual longitudinal feature as a discrete token. The transpose of the input matrix 𝒳^*t*^ ∈ ℝ^*C×T*^ is viewed as a collection of *C* tokens, each of dimension *T*. This approach allows the self-attention mechanism to model the cross-correlations between specific genomic alterations and panel-level dynamics across the entire treatment window. By focusing on the “feature” dimension, the network learns to weight the relative importance of specific gene-level trajectories (e.g., *TP* 53 dynamics vs. *KRAS* dynamics) in determining patient risk. To calculate the Cox proportional hazards log-risk, the attention-weighted representations are concatenated with frozen pre-trained embeddings and clinical variables, then passed through a fully connected layer, using batch normalization prior to concatenation and followed by a dropout layer with a regularizing probability of 0.4—consistent with all competing neural network frameworks.

### Recurrent Neural Network/Gated Recurrent Unit Architectures

We implemented temporal encoding branches based on recurrent architectures to capture sequential dependencies in longitudinal ctDNA data. We evaluated both a *Recurrent Neural Network* (RNN) and a *Gated Recurrent Unit* (GRU) configuration. To focus on the longitudinal evolution of the patient’s molecular profile, these models utilize the input data, 𝒳 ∈ ℝ^*T ×C*^, where each input token at index *t* represents the complete vector of 21 cfDNA/ctDNA features (17 gene-level VAFs and 4 panel-level features) at that specific clinical time point. The recurrent models process the sequence {*x*_1_, *x*_2_, *x*_3_, *x*_4_}iteratively, updating a hidden state *h*_*t*_ at each timestep to maintain a memory of the preceding therapy cycles. For the GRU variant, gating mechanisms (reset and update gates) are employed to regulate the flow of information and mitigate the vanishing gradient problem common in standard RNNs. To obtain a fixed-length representation of the entire treatment trajectory, we extract the latent feature vector from the final timestep (*T* = 4). This terminal hidden state serves as a comprehensive summary of the longitudinal ctDNA dynamics up to the *C*4*D*1 assessment. The final stage of the recurrent framework follows a fusion strategy identical to every competing models. The latent temporal representation is concatenated with the frozen embeddings *e*_*t*_ from the pre-trained genomic-interaction encoder and the baseline clinical characteristics *c*, using batch normalization prior to concatenation and followed by a dropout layer with a regularizing probability of 0.4—consistent with all competing neural network frameworks. The parameters of the RNN/GRU modules are optimized jointly with the output layer by minimizing the negative log-partial likelihood of the Cox proportional hazards model.

### Multilayer Perceptron (DeepSurv) Architecture

For a non-sequential baseline, we implemented a *Multilayer Perceptron (DeepSurv)* configuration. The Multilayer Perceptron processes the input 𝒳 ∈ℝ^*T ×C*^ as a flattened vector of all longitudinal features. To perform dimensionality reduction and capture high-level feature interactions, the input is passed through a fully connected layer where the hidden dimension is reduced by half relative to the input size. This bottleneck representation is activated via a sigmoid function to introduce non-linearity.

The resulting compressed feature vector is then integrated with the frozen pre-trained per-timestep genomic-interaction embeddings *e*_*t*_ and baseline clinical characteristics *c* to form the unified feature vector Φ, using batch normalization prior to concatenation and followed by a dropout layer with a regularizing probability of 0.4—consistent with all competing neural network frameworks. As with the other architectures, Φ is mapped to the predicted log-risk *η* for optimization via the partial likelihood of the Cox proportional hazards model.

### Functional Principal Component Random Forest (FPCRF)

As a baseline for trajectory-based modeling, we adapted the *Functional Principal Component Random Forest* (FPCRF) framework proposed by Ding et al. [12]. In this workflow, longitudinal ctDNA and cfDNA data are treated as functional objects rather than discrete sequences. For each of the 21 longitudinal features—comprising 17 gene-level VAFs and 4 panel-level features—we extracted two Functional Principal Components (FPCs) to capture the underlying continuous temporal variations, resulting in a total of 42 extracted features.

As explained in Ding et al. [12], to manage the high-dimensional nature of the extracted FPCs, we utilized a Random Forest (RF) variable importance measure to rank and select the 20 most predictive features among the 42 FPC-derived features. To further mitigate collinearity and condense the feature space, standard Principal Component Analysis (PCA) was applied to the top-ranked FPCs [12], reducing the dimensionality to 10 principal components.

While the original FPCRF method was designed for a specific set of 17 panel-level features, we modified the input space to include the 21 longitudinal features used across all competing models in this study. Furthermore, to ensure a fair comparison with our SHER-LOC and all competing methods, we extended the original FPCRF framework by integrating the FPC-derived representations with the frozen pre-trained per-timestep genomic-interaction embeddings *e*_*t*_ and baseline clinical characteristics *c* inside a Cox proportional hazards with Elastic Net penalty as described in Ding et al [12]. By incorporating the pre-trained embeddings *e*_*t*_—a component not present in the original implementation—we ensured that performance differences across models were attributable to their temporal encoding strategies rather than disparities in available information.

### Random Survival Forest (RSF)

As a non-parametric machine learning baseline, we implemented the *Random Survival Forest* (RSF) algorithm as described by Ishwaran et al. [34]. RSF is an ensemble tree-based method that extends Breiman’s random forest framework to right-censored survival data by growing *B* trees. At each node of a tree, a random subset of variables is selected, and a splitting criterion—the log-rank test—is used to maximize the difference in survival between daughter nodes. The prediction mechanism of RSF relies on the cumulative hazard function (CHF) calculated at the terminal nodes. For any given tree, let *h* be a terminal node. The CHF estimate for all cases falling within *h* is calculated using the Nelson–Aalen estimator:

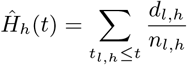

where *d*_*l,h*_ and *n*_*l,h*_ are the number of deaths and individuals at risk at time *t*_*l,h*_ within node *h*, respectively. Consequently, all cases within the same terminal node *h* are assigned the same CHF. These individual tree estimates are then averaged across the entire forest to produce the ensemble CHF [34].

For this implementation, the input data was structured by flattening the 21 longitudinal features (17 gene-level VAFs and 4 panel-level features) into a single high-dimensional vector. To ensure a fair comparison with all competing methods, this flattened temporal representation was concatenated with the frozen pre-trained per-timestep genomic-interaction embeddings *e*_*t*_ and baseline clinical characteristics *c* to form the unified feature vector Φ. Tuning for the Random Survival Forest was performed on the training data over a grid comprising 50 and 100 estimators combined with a minimum sample split threshold of 2 and 6.

### Joint Model Architecture

To provide a robust statistical baseline, we implemented a multivariate *Joint Model* (JM) using the JMbayes2 framework in R [40]. This approach simultaneously models the longitudinal trajectories of gene- and panel-level cfDNA features. To ensure numerical stability within the joint models, we only included features with a sparsity of less than 95% (i.e., those with non-zero values in at least 5% of the training set).

#### Sub-model Specifications

The longitudinal process for each cfDNA feature was defined by a linear mixed-effects (LME) submodel as a function of time, utilizing a fixed linear slope and a patient-specific random intercept:

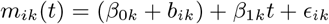

where *b*_*ik*_ represents the random intercept for feature *k* from patient *i*. This configuration assumes that baseline levels of ctDNA vary across individuals while the rate of change per treatment cycle follows a common population trend.

The longitudinal sub-models are linked to the survival sub-model via a shared-parameter framework. The hazard function for patient *i* is defined as follows:

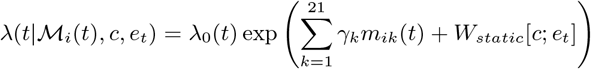

Here, *λ*_0_(*t*) denotes the baseline hazard,ℳ _*i*_(*t*) = {*m*_*i*1_(*t*), …, *m*_*ip*_(*t*)} represents the set of current values for the *p* longitudinal cfDNA markers at time *t*. The model incorporates frozen pre-trained genomic-interaction embeddings (*e*_*t*_) and baseline clinical characteristics (*c*), with *γ*_*k*_ capturing the association parameters for each longitudinal trajectory.

#### Bayesian Estimation and MCMC Parameters

The parameters of the longitudinal and survival sub-models were estimated simultaneously within a Bayesian framework using Markov Chain Monte Carlo (MCMC) sampling. This allows the model to link the true unobserved longitudinal trajectory *m*_*ik*_(*t*) to the hazard of the event through association parameters. To ensure MCMC convergence in `JMbayes2`, we maintained the default three parallel chains while increasing the total iterations to 4,000 and the burn-in period to 1,000 (standard defaults are 3,500 and 500, respectively).

### Ridge, Lasso and Elastic Net Architectures

We trained penalized Cox proportional hazards models by minimizing the negative log-partial likelihood with an added regularization term. For patient *i*, the linear predictor was defined as *η*_*i*_ = *X*_*i*_*β*, where *X*_*i*_ denotes the covariate vector and *β* the regression coefficients. The objective function was

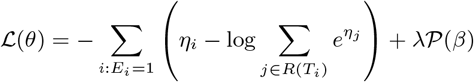

where *E*_*i*_ is the event indicator and *R*(*T*_*i*_) is the risk set at time *T*_*i*_. The penalty term *𝒫* (*β*) corresponds to *Ridge* 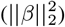, *Lasso* 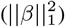,or *Elastic Net* 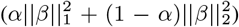 regularization, where the mixing parameter was fixed to *α* = 0.5. The hyperparameter *λ*, controlling the overall strength of regularization, was selected via 4-fold cross-validation on the training set. *Ridge, Lasso*, and *Elastic Net* Cox models process the input 𝒳∈ ℝ^*T ×C*^ as a flattened vector of all longitudinal features (17 gene-level VAFs and 4 panel-level features), concatenated with pre-trained genomic-interaction embeddings *e*_*t*_ and baseline clinical characteristics *c*.

### Statistical Evaluation and Model Robustness

In Figure 2 and Supplementary Figure 1, the SHERLOC ctDNA-based risk score was trained exclusively on ctDNA samples collected from baseline to C3D1, to ensure a fair comparison with RECIST assessment, which was also performed at C3D1. After excluding patients who experienced an OS event prior to the C3D1 landmark, 398 patients were retained. Among these, 395 had available C3D1 RECIST assessments. To evaluate the independent prognostic value of the SHERLOC ctDNA-based risk score, a random 70/30 train–test split was applied (n = 276 and n = 119 patients, respectively). The Kaplan–Meier curves and forest plots shown in Figure 2 present results for the independent test set.

To ensure the stability and reproducibility of our benchmarks (Tables 1, 2, 3), we employed a repeated train-test split approach consisting of 100 repeats (Monte Carlo cross-validation). For each experience, the dataset was partitioned using a unique random seed into a training set (70%) and a test set (30%). Models were fit on the training data and evaluated on the corresponding test data, with performance reported as the mean across all 100 test sets. To account for the statistical dependency introduced by overlapping training data across repetitions, we calculated a corrected standard deviation using the formula for resampled estimates, following the methodology of Nadeau and Bengio [25]:

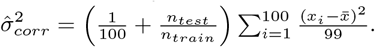

This correction provides a more rigorous measure of model variability [25].

### Statistical Evaluation of Model Representations

The SHERLOC architecture enables fine-grained decomposition of prognostic information by integrating representations from the ctDNA-dynamics encoder, the genomic-interaction encoder, and clinical covariates into a single fully connected Cox proportional hazards log-risk layer. To quantify the individual contribution of each learned representation, we adopted a resampling-based validation framework in which all upstream encoders were frozen and only the final Cox layer was retrained. This procedure was repeated across (N = 100) resampled splits, yielding an empirical distribution of coefficients for each representation. Statistical significance was assessed using the corrected resampled *t*-test of Nadeau and Bengio [25], which accounts for dependencies induced by overlapping training sets in resampling-based evaluation. The corrected variance estimate was used to compute standard deviations, confidence intervals, and *P* -values, enabling valid inference under the non-independent sampling scheme.

To evaluate gene-level contributions across the full architecture, feature importance was examined within both the longitudinal ctDNA-dynamics encoder and the pre-trained genomic-interaction encoder. Because the same gene-level VAF is processed through two distinct architectural arms with different latent regularization schemes, direct interpretation of raw coefficients *β* can be confounded by opposing signs. To address this, we report feature importance as the mean of the coefficient distribution in absolute value, emphasizing the magnitude of association with survival rather than directionality. Critically, this transformation is applied only to the point estimate of the mean, while the original standard deviation of the raw coefficient distribution is strictly preserved for all downstream inference.

### Performance Metrics for Survival Analysis

Model performance was evaluated using time-dependent metrics adjusted via Inverse Probability of Censoring Weighting (IPCW). We computed the Cumulative/Dynamic AUC to assess performance in terms of survival discrimination. Let *y*_*i*_ denote the observed time for patient *i*. For a given time *t*, this metric quantifies the model’s ability to distinguish subjects who experience an event by time *t* (cases) from those who do not (controls), defined as:

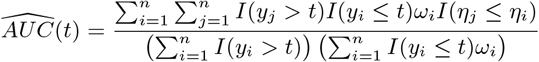

where *ω*_*i*_ are IPCW weights and *η* is the predicted risk. The integrated AUC (*iAUC*) provides a summary measure over the time interval [*τ*_1_, *τ*_2_] (in our analysis from the 3rd month to the 30th month) weighted by the survival function *Ŝ*(*t*):

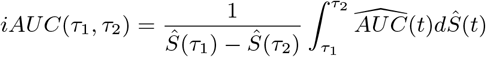

To evaluate the discriminative power of models, we calculated the concordance index (*C*-index) [28]. Unlike traditional measures, this estimator incorporates Inverse Probability of Censoring Weighting to provide a consistent estimate of model accuracy that is robust to heavy censoring. Specifically, for a truncation time *τ*, the index is calculated as:

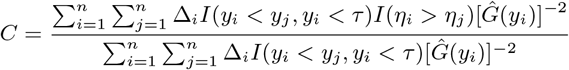

where *y*_*i*_ represents the observed survival time, Δ_*i*_ is the event indicator, *η*_*i*_ is the predicted risk score, and 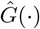 is the Kaplan-Meier estimator of the censoring distribution. This approach ensures that the evaluation remains valid across the follow-up period by weighting the informative pairs by the inverse of the probability of being censored. In our analysis, the truncation time *τ* for the computation of the *C*-index was set to 30 months.

Finally, we employed the Integrated Brier Score (IBS) to evaluate models’ mean squared prediction error (using scikit-survival python library). The time-dependent Brier Score at time *t* measures the weighted mean squared error between the predicted survival probability *Ŝ*(*t*|*x*) and the actual status:

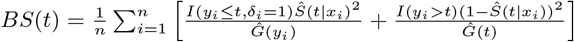

where 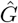represents the Kaplan-Meier estimate of the censoring distribution. The IBS integrates this score over the specified range, estimated via the trapezoidal rule:

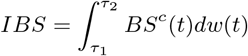

with weighting *w*(*t*) = *t/*(*τ*_2_ *−τ*_1_), where a score of 0 represents perfect prediction. In our analysis, the brier score is integrated from the 3rd month to the 30th month, matching the integration period used for the AUC.

## Acknowledgements

A. Mamann, H. Benkirane, E. Bernard, B. Besse, P.H. Cournè de and S. Michiels were supported by Prism—National Precision Medicine Center in Oncology funded by the France 2030 program and the French National Research Agency (ANR) [grant number ANR-23-IAHU-0002]. A. Mamann and B. Besse were supported by RHU Reveal funded by the France 2030 programme and the French National Research Agency (ANR) [grant number REVEAL ANR-21-RHUS-0013]. E. Bernard was supported by the INSERM ATIP-Avenir Program.

## Author contributions

A. Mamann is the first author, and was the main contributor to every part of the study. A. Mamann, J. Das, S. Michiels and P.H. Cournè de contributed on the conception of SHERLOC. A. Mamann, H. Benkirane, J. Das, F. Bugiotti, S. Michiels and P.H. Cournè de contributed on the implementation of competing models in the benchmark. A. Mamann, E. Bernard, B. Besse, S. Michiels and P.H. Cournè de contributed on the clinical applications of SHERLOC. A. Mamann, S. Michiels and P.H. Cournè de drafted the manuscript. A. Mamann, J. Das, H. Benkirane, F. Bugiotti, E. Bernard, B. Besse, S. Michiels and P.H. Cournè de performed critical reviews. All authors discussed the results, contributed to the final work, and have provided final approval of the completed version.

## Competing interests

S. Michiels reports, outside the scope of the submitted work, fees for scientific committee study member for Roche; and data and safety monitoring member for clinical trials for IQVIA, Kedrion, Biophytis, Servier, and Yuhan. B. Besse reports, outside the scope of the submitted work, personal fees from AbbVie, AstraZeneca, Biontech SE, Beijing Avistone Biotechnology, Bristol Myer Sqibb, Gilead, GSK, MSD, Pharmamar, Regeneron, Roche Diagnostics, Sanofi Aventis, Servier, CureVac AG, Eli Lilly, F.Hoffmann–La Roche Ltd, Foghorn Therapeutics Inc, OncoC4 Inc, Tubulis GmbH Conseil, Amgen, Beigene, GENMAB A/S, Ose Immunotherapeutics, Helios Medical Communications Ltd, Ose Immunotherapeutics, and Takeda during the conduct of the study. No other disclosures were reported. All remaining authors have declared no conflicts of interest.

## Declaration of generative AI and AI-assisted technologies in the writing process

During the preparation of this work, the author(s) used Gemini 3 in order to refine the grammatical structure and improve the readability of the manuscript text. After using this tool/service, the author(s) reviewed and edited the content as needed and take(s) full responsibility for the content of the published article.

## Data availability

All clinical and ctDNA data for **IMpower150** are deposited to the European Genome-Phenome Archive under accession number **EGAS00001006703**. Qualified researchers may request access to individual patient-level data through the clinical study data request platform (https://vivli.org/). Further details on Roche’s criteria for eligible studies are available at https://vivli.org/members/ourmembers. For further details on Roche’s Global Policy on the Sharing of Clinical Information and how to request access to related clinical study documents, see https://www.roche.com/research_and_development/who_we_are_how_we_work/clinical_trials/our_commitment_to_data_sharing.htm.

All clinical and DNA sequencing data for the **MSK-CHORD** cohort are publicly accessible via cBioPortal under the study identifier **msk chord 2024** (https://www.cbioportal.org/study/summary?id=msk_chord_2024). The dataset is made available under a Creative Commons Attribution-NonCommercial-NoDerivs 4.0 International (CC BY-NC-ND 4.0) license. Because this license restricts commercial utilization, qualified researchers or entities seeking commercial access or specialized licensing terms may submit inquiries to the Memorial Sloan Kettering Cancer Center (MSK) Data Requests team. Researchers utilizing these data must cite the original publication (*MSK-CHORD, Nature 2024*) [24].

## Code availability

SHERLOC is fully open source. The complete source code, including preprocessing pipelines, encoder implementations, and training scripts for SHERLOC and all competing models included in the benchmark, is publicly available. The repository has been designed to facilitate reproducibility and ease of use, enabling the community to apply, evaluate, and build upon the proposed framework.

- **Repository:** https://github.com/AaronMamannToledano/SHERLOC

---

**Supplementary Figure 1:**
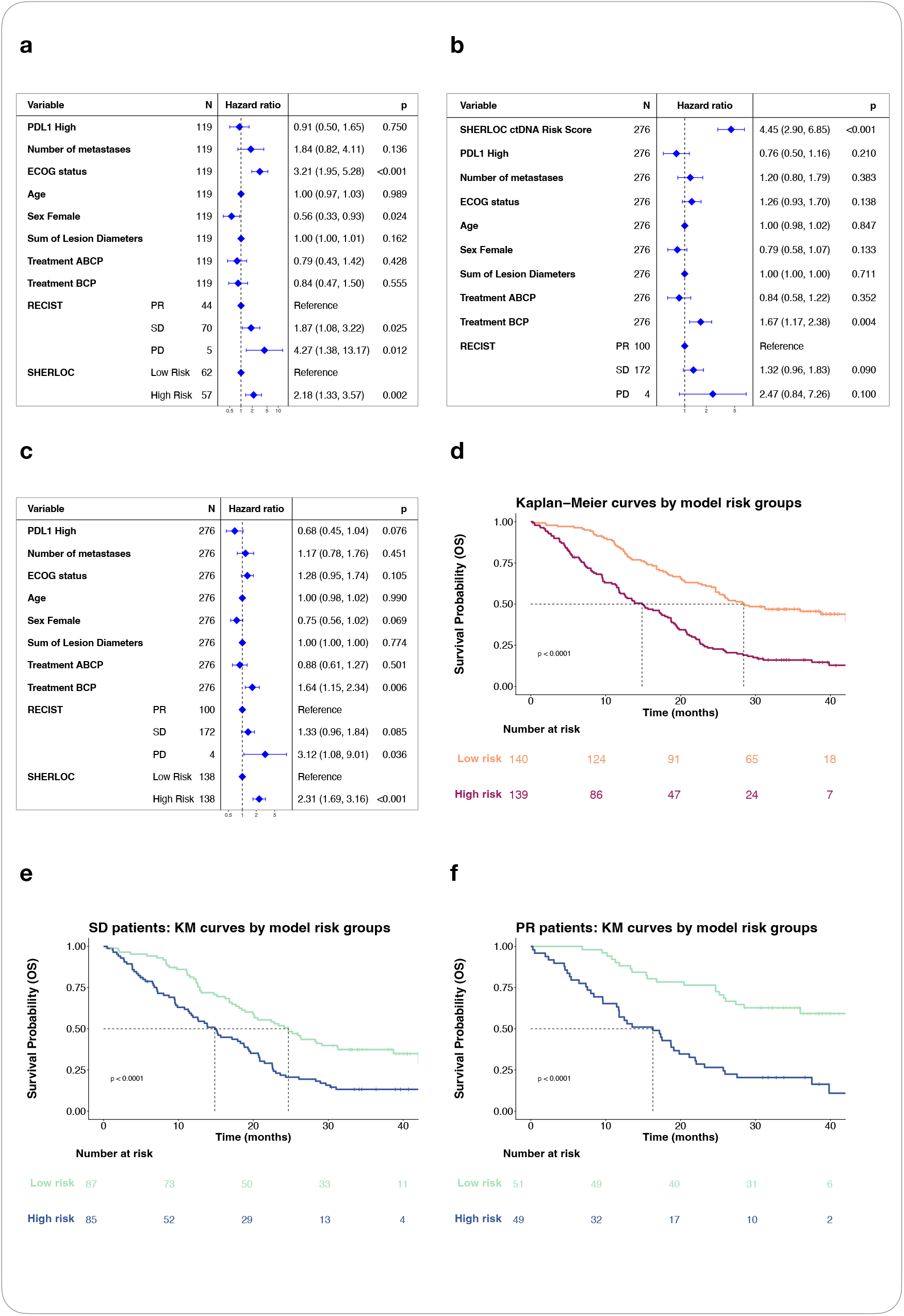
SHERLOC ctDNA-based risk model improves upon traditional RECIST criteria. **(a)** OS hazard ratio (HR) of a multivariable Cox regression in the test set (*n* = 119) incorporating clinical features, RECIST categories assessed at C3D1 and ctDNA-based model risk categories (Low/High Risk) derived from SHERLOC trained only on ctDNA samples collected from baseline to C3D1. OS was recalculated from the RECIST assessment date (landmark analysis). Of 398 patients remaining after pre-landmark OS exclusion, 395 had C3D1 RECIST assessments. **(b, c)** OS hazard ratio (HR) of a multivariable Cox regression in the training set (*n* = 276) incorporating clinical features, RECIST categories assessed at C3D1 and ctDNA-based model risk score **(b)** or risk categories **(c)** derived from SHERLOC trained only on ctDNA samples collected from baseline to C3D1. **(d)** OS Kaplan-Meier curves in the training set stratified by the SHERLOC ctDNA-based risk categories. Median risk scores of the training set were used as a cut point to derive low- and high-risk patients. **(e, f)** OS Kaplan-Meier curves in the training set for patients categorized as SD **(e)** or PR **(f)** from RECIST and stratified by the ctDNA-based model risk categories. **Abbreviations:** CR, complete response; PD, progressive disease; PR, partial response; SD, stable disease.

